# Quantifying Three-dimensional Chromatin Organization Utilizing Scanning Transmission Electron Microscopy: ChromSTEM

**DOI:** 10.1101/636209

**Authors:** Yue Li, Eric Roth, Vasundhara Agrawal, Adam Eshein, Jane Fredrick, Luay Almassalha, Anne Shim, Reiner Bleher, Vinayak P. Dravid, Vadim Backman

**Affiliations:** Applied Physics Program, Northwestern University, Evanston, Illinois 60208, USA; Department of Materials Sciences and Engineering, Northwestern University, Evanston, Illinois 60208, USA; Department of Biomedical Engineering, Northwestern University, Evanston, Illinois 60208, USA

## Abstract

Chromatin organization over a wide range of length scales plays a critical role in the regulation of gene expression and deciphering these processes requires high-resolution, three-dimensional, quantitative imaging of chromatin structure *in vitro*. Herein we introduce ChromSTEM, a method which utilizes high angle annular dark field imaging and tomography in scanning transmission electron microscopy in combination with DNA-specific staining for electron microscopy. We utilized ChromSTEM to quantify chromatin structure in cultured cells and tissue biopsies through local DNA distribution and the scaling behavior of chromatin polymer. We observed that chromatin is densely packed with an average volume concentration of over 30% with heterochromatin having a two-fold higher density compared to euchromatin. Chromatin was arranged into spatially well-defined nanoscale packing domains with fractal internal structure and genomic size between 100 and 400 kb, comparable to that of topologically associated domains. The packing domains varied in DNA concentration and fractal dimension and had one of the distinct states of chromatin packing with differential ratio of DNA content to the chromatin volume concentration. Finally, we observed a significant intercellular heterogeneity of chromatin organization even within a genetically uniform cell population, which demonstrates the imperative for high-throughput characterization of chromatin structure at the single cell level.

## Introduction

Regulation of gene transcription is essential in sustaining normal cell function, controlling cell differentiation and determining cell fate, and transcriptional alterations have been implicated in a variety of diseases including cancer and cardiovascular, developmental, neurological, and autoimmune disorders ^1, 2, 3^. While early studies, which focused on the molecular mechanisms of transcriptional regulation based on a linear model of genome organization, provided significant knowledge of the genome regulation, they also demonstrated the substantial limitations of this approximation ^4, 5, 6, 7, 8, 9, 10^. Recent studies have unambiguously shown that genes can interact with multiple distal elements within distances up to several Mb away, suggesting a machinery of transcriptional regulation based on the three-dimensional (3D) chromatin structure ^11, 12, 13, 14, 15^. While the genetic information is encoded in the linear sequence, the appropriate gene transcription requires complex 3D genome organization ^16^.

A number of methods have been developed to analyze 3D chromatin structure, including chromatin conformation capture (e.g. Hi-C), neutron scattering, soft x-ray tomography (SXT), and super-resolution microscopy ^17, 18, 19, 20, 21, 22^. These techniques have provided critical insights into the principles of 3D chromatin structure. However, they have fundamental limitations such as the inability to quantify the spatial distribution of chromatin (C-based, neutron scattering) or limited resolution (30-50 nm for SXT, super-resolution microscopy) ^23^. To precisely image chromatin down to the level of a single nucleosome (11nm) and DNA strands (2nm) and map the 3D chromatin architecture within the nucleus, electron microscopy (EM) remains the technique of choice.

Recently, Ou et al. reported a new transformative approach, which utilizes “click-EM” staining that specifically labels DNA and multi-axis transmission electron microscopy (TEM) tomography, for chromatin imaging with a nominal resolution of 1.6 nm in 3D (ChromEMT) ^21, 24^. Using ChromEMT Ou et. al. demonstrated that chromatin is a disordered 5 to 24 nm in diameter polymer chain, rather than the classically considered hierarchically folded assembly ^24^. As TEM imaging contrast is non-linear (confounded by phase, diffraction, and mass-thickness contrast), the application of TEM-based imaging to quantify chromatin packing in terms of physical mass-density distribution is challenging. Furthermore, it is difficult to perform ChromEMT for the whole cell, as it would require dozens of serial-sections to cover a mammalian nucleus (~6 μm in height). The challenge becomes ever more daunting when a comparative analysis of several cells or cell types exposed to differential conditions is required. The complexity is further exacerbated by the phenomenon of genomic and transcriptional intercellular heterogeneity, which most cell populations exhibit, and which would necessitate the analysis of an ensemble of cells.

To develop a quantitative and high-throughput method for high-resolution, 3D chromatin imaging, we adapted the ChromEMT framework to incorporate STEM high angle annular dark field (STEM HAADF) imaging. This hybrid method, ChromSTEM, allows quantitative imaging through Z-contrast, which originates from Rutherford scattering by *I*~*Z*^1.7^, where *I* is the image contrast, and *Z* is the atomic number of the atom on the electron trajectory ^25^. In the case of ChromSTEM, the osmium bound to the chromatin dominates the image contrast, and the observed image intensity is proportional to the DNA mass-density. We demonstrate the utility of ChromSTEM by reconstructing the 3D chromatin structure of adenocarcinoma human alveolar basal epithelial (A549) cells at a nominal voxel resolution of 3 nm. We combined the 3D data with ultra-thin section imaging and developed a statistical method to accurately quantify chromatin packing through DNA density distribution, chromatin volume concentration (CVC), chromatin polymer mass-scaling, and nuclear compartment positioning for the whole nucleus. We quantified the packing properties of the euchromatin and heterochromatin compartments. We also observed that chromatin is organized into spatially separated 100 - 200 nm packing domains, with fractal internal structure and genomic size comparable to topologically associated domains (TADs). Across the genome, packing domains varied significantly in their fractal dimension, DNA content, CVC, and packing density heterogeneity.

## Results

### ChromSTEM platform for analysis of chromatin organization

3D chromatin organization governs gene transcription by controlling genome connectivity, DNA accessibility, and transcriptional heterogeneity ^26, 27, 28^. Emerging evidence shows that the proper positioning of the chromatin within the nucleus also plays an indispensable role in maintaining normal transcriptional function ^10, 29^. Utilizing the ChromSTEM platform, one can obtain quantitative information on the native 3D chromatin architecture. Fig. 1 shows the roadmap towards a comprehensive analysis of the chromatin organization.

**Figure 1.**
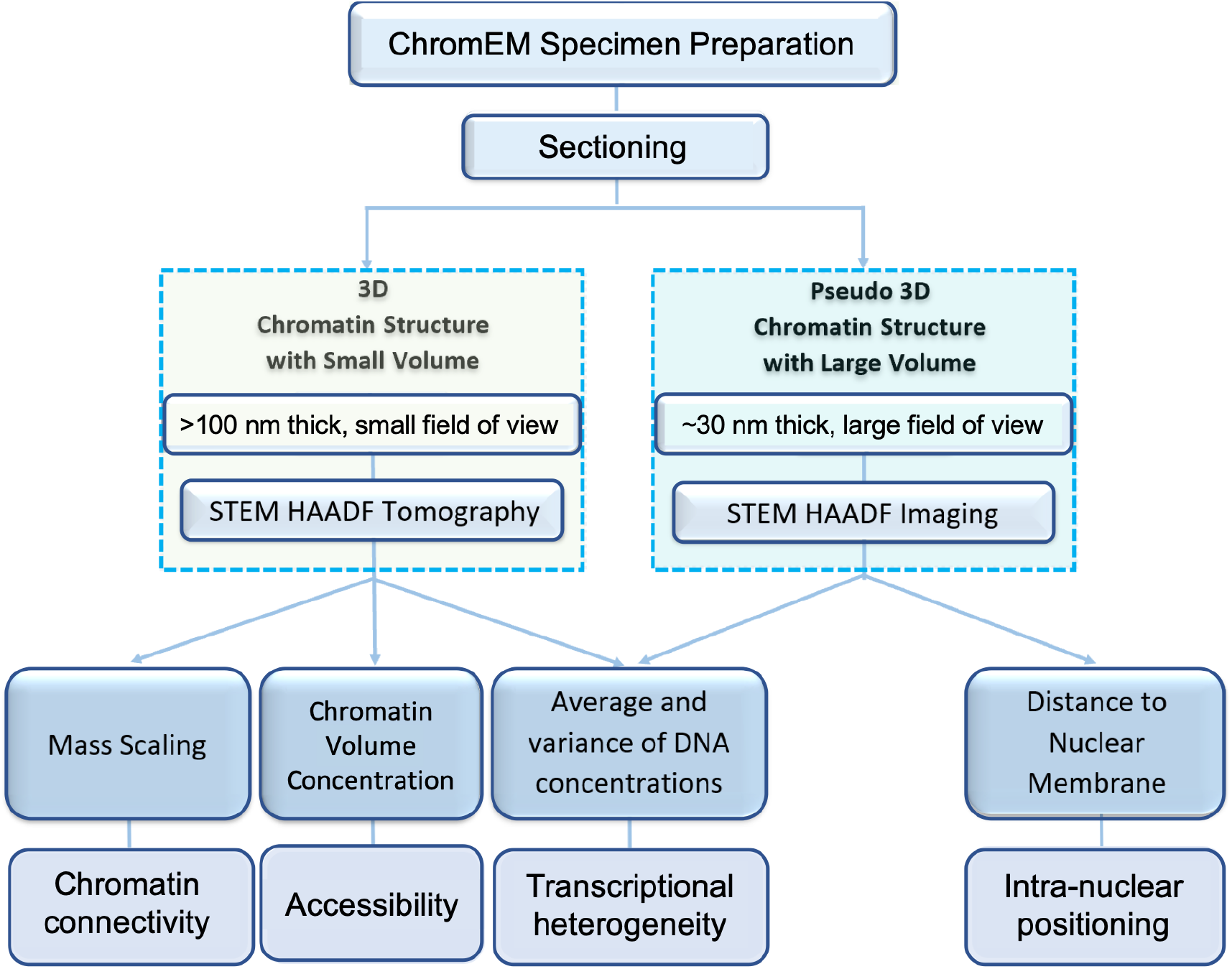
Multi-scale quantification of the chromatin organization through ChromSTEM. The ChromSTEM platform includes several experimental techniques to assess chromatin 3D organization by directly measuring the chromatin mass scaling, chromatin volume concentration (CVC), the average and the variance of DNA concentration, the mass scaling exponent (D), and the distance to the nuclear membrane. To estimate chromatin connectivity and accessibility, 3D tomography reconstruction of semi-thick sections (~>100 nm thickness) can be deployed. The analysis of the spatial distribution of the chromatin throughout the nucleus is achieved by ChromSTEM mapping of a stack of ultrathin sections (~ 30 nm thickness). Both tomography and thin section projection can be employed to measure chromatin heterogeneity.

Following the ChromEM protocol reported previously, we labeled the DNA of A549 cells, and fluorescence images were acquired (Fig. 2a, b) during photo-oxidation. After resin embedding, the labeled regions can be identified based on image contrast in bright field optical micrographs: the photo-oxidized cells appeared significantly darker than the non-photobleached cells (Fig. 2c). Dual-tilt STEM tomography in HAADF mode was performed for part of the nucleus containing a hetero/euchromatin interface on a 100 nm resin section (Fig. 2d). We observed continuous variations of the image contrast inside the nucleus, different from the near binary image contrast from conventional EM staining method. Each tilt series was aligned with fiducial markers in IMOD (Supplementary, Mov. 1, 2) and reconstructed by a penalized maximum likelihood (PLM-hybrid) algorithm in Tomopy ^30, 31^. The two sets of tomograms were combined in IMOD to suppress missing cone artifacts (Supplementary Fig. 1)^32^. The final tomography (Fig. 2f) has a nominal voxel size of 2.9 nm (Supplementary, Mov. 3), with clearly resolved nucleosomes (Fig.2g) and linker DNA (Fig. 2h). We also identified several distinct higher order supranucleosomal structures such as stacks and rings (Fig. 2i, j). We rendered the 3D volume of the chromatin in the volume viewer in FIJI (Fig. 2k, l, Supplementary, Mov. 4,5,6). The voxel intensity of the tomogram was used for color-coding ^33^. The chromatin chain was comprised of 11 - 18 nm “core” regions with high DNA density (Fig. 2l, pink) surrounded by the 3 - 8 nm “shell” with low DNA density (green). The total diameter of the chromatin chain (core and shell) ranged from 14 to 26 nm, supporting the size of the fibrous polymer previously identified using ChromEMT^24^.

**Figure 2.**
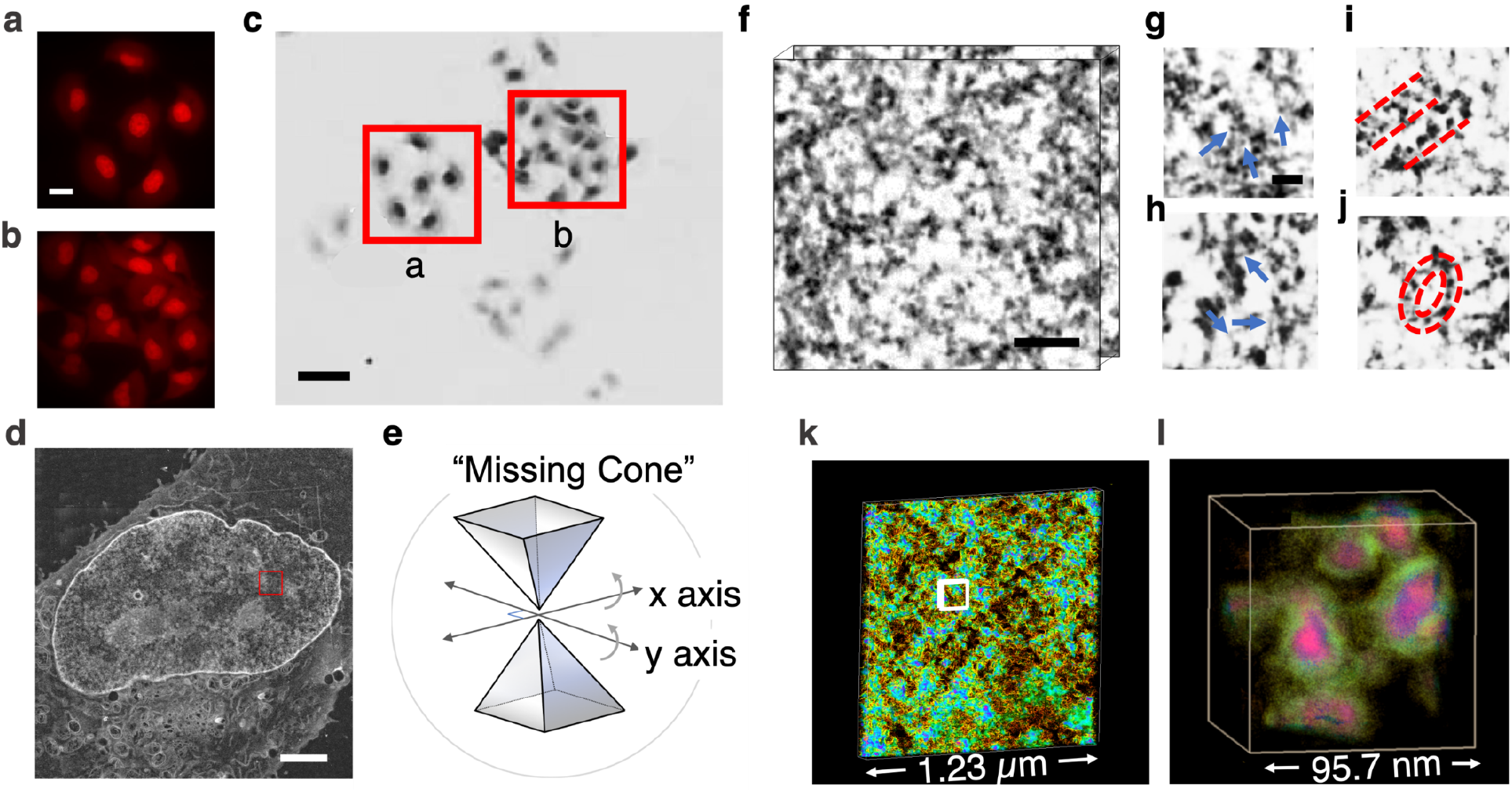
ChromSTEM tomography reconstruction of the chromatin of an A549 cell. **(a-b)** The DRAQ5 photo-oxidation process takes 7 min for each region of interest. Scale bar: 10 μm. **(c)** The labeled regions were more intensely stained than the nearby regions (red squares, the letter corresponds to the regions in the left panels). Scale bar: 20 μm. **(d)** STEM image of a 100 nm thick section of an A549 cell in HAADF mode. Scale bar: 2 μm. **(e)** Schematics for dual-tilt tomography. The sample was tilted from −60° to 60° with 2° increments on two perpendicular axes. **(f)** 3D tomography of the A549 chromatin. Scale bar: 120 nm. **(g-j)** The fine structure of the chromatin chain: Nucleosomes (blue arrows in **(g)**), linker DNA (blue arrows in **(h)**) supranucleososomal stack (red dashed lines in **(i)**) and ring (red dashed circles in **(j)**). Scale bar: 30nm. **(k-l)** 3D rendering of the chromatin organization, the pseudo color was based on the intensity of the tomograms (Supplementary Mov. 4-6). **(I)** A magnified view of the region labeled by a white square in **(k)**. In **(I)**, pink and green represent high and low DNA density regions, respectively.

### Chromatin internal packing structure

Topologically associated domains (TADs) are functionally defined, sub-Mbp scale genome structures that might be comprised of several chromatin loops ^34^. TADs are believed to play a critical role in gene transcription by increasing interactions between loci located within the same domain and, equally importantly, insulating genes from genomic regions located in neighboring domains ^35^. Whether TADs are physical elements of chromatin packing or statistics of an ensemble of chromatin states across a cell population has been a subject of an ongoing debate ^36, 37, 38^. While most experimental HiC data have been acquired at the cell population level (with the exception of lower resolution single-cell HiC) ^39, 40, 41^, TAD-like nanocompartments and nanoclusters have been reported with super-resolution techniques at the single cell level ^21, 42, 43, 44^. Given its high spatial resolution and the ability to image a population of cells, ChromSTEM has the potential to address some of these controversies.

Fig. 3a shows a HAADF image of an A549 cell nucleus, which indicates that the chromatin organized into multiple packing domains (insets). To characterize the domain structure, we performed the mass-scaling analysis. Similar to other polymeric systems, the scaling behavior of chromatin describes the relationship between the physical size (radius *r*) of a chromatin region and the genomic length (chromatin mass or the number of base pairs, *M*) contained within it: *M* ∝ *r*^*D*^, where *D* is the chromatin packing density scaling or the fractal dimension of a given chromatin domain. *D* of an unconstrained free polymer in equilibrium may range from *D* = 5/3 (for an excluded volume polymer) to *D* = 3 (for space-filling polymer) depending on the balance between the free-energy of polymer-polymer and polymer-solvent interactions. *D* is further modulated by constrain processes, such as chromatin loops. If a polymer forms a number of spatially uncorrelated domains, the mass-scaling of the supradomain structure (i.e. *r* is greater than the domain size) is also 3 but the structure is no longer fractal at these length scales.

**Figure 3.**
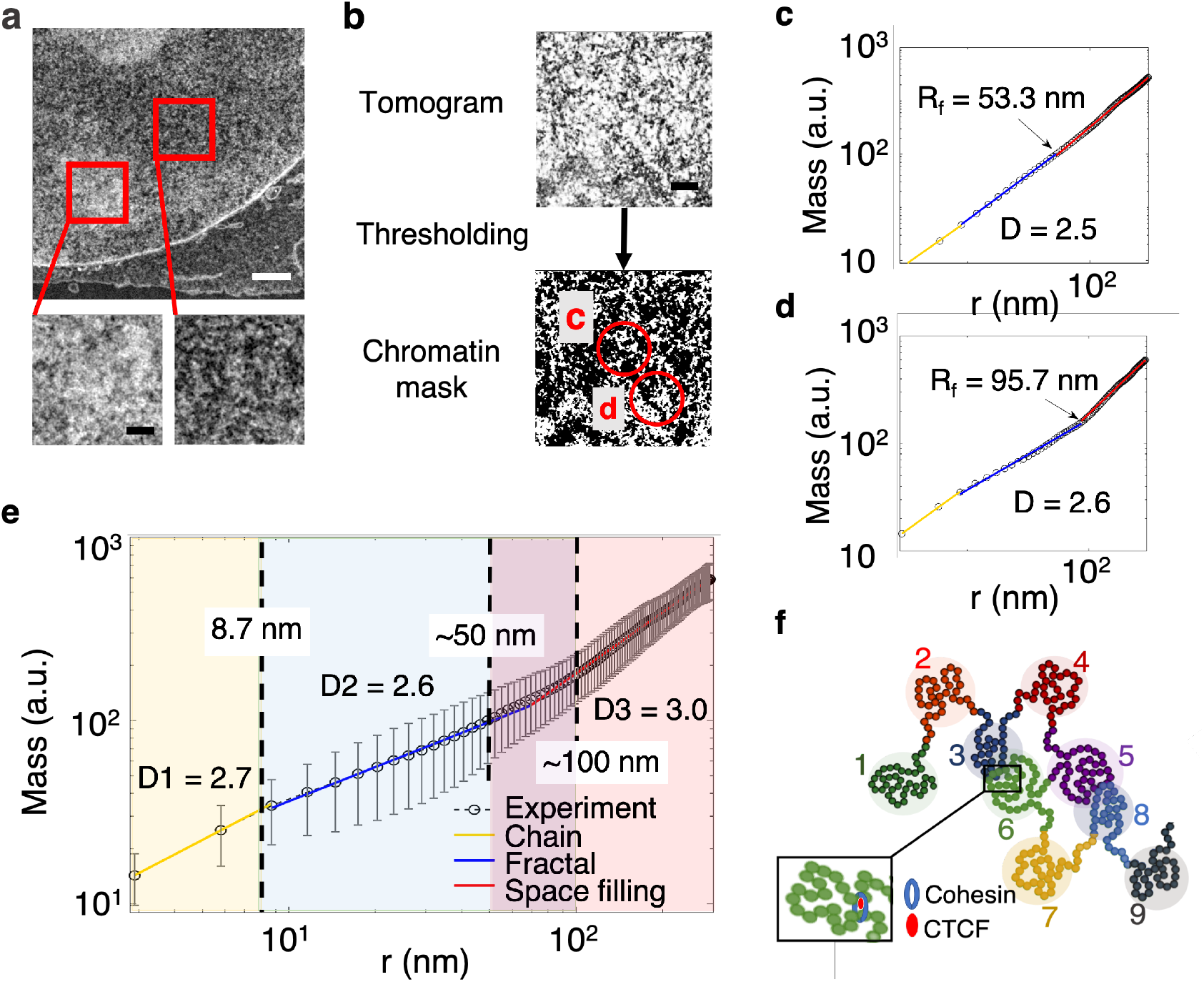
The internal structure of chromatin packing domains. **(a)** STEM HAADF image of an 100 nm section of an A549 cell nucleus shows that the chromatin is organized into compartments (light and dark image contrast), which consist of smaller domains (insets). Scale bar: 1 μm; inset scale bar: 300 nm. **(b)** For each virtual 2D slice, automatic thresholding was applied to create a binary chromatin mask. The mass scaling analysis was performed inside a circle with a radius of 300 nm. Scale bar: 120 nm. **(c-d)** Examples of the mass scaling curve for chromatin regions highlighted by red circles c and d in **(b)**. In (c-d), Mass is defined as the mass of chromatin that is enclosed by a ring with inner radius r and the width of 2.9 nm (1 pixel). Three distinct regimes of mass scaling can be identified: the chromatin chain regime (yellow data points and regression line), the packing domain with a fractal internal structure (blue), and the supradomain structure (red). The fractal dimension (D) and the radial size (*R*_*f*_, defined as the upper length scale of fractality) of domains varied across domains. **(e)** The average mass scaling of 4096 randomly centered regions on each of the 33 tomograms. The error bar is the standard deviation. Three regimes of mass-scaling behavior can be identified: the chromatin chain regime (yellow) with r < 8.7 nm, fractal regime (blue) with the average fractal dimension D = 2.6 and 8.7 nm < r < 50nm, space-filling regime of spatially uncorrelated packing domains (red) with scaling exponent = 3 and r > 100 nm. In between the fractal region and space-filling region ranging from 50 nm to 100 nm (purple), we observed a smooth increase of the scaling exponent, which indicates a variability in packing domains sizes. **(f)** Schematic of the packing structure of chromatin that is suggested by the ChromSTEM data. The schematic shows 9 chromatin fractal packing domains, some of which might be formed by loop extrusion (inset) or non-loop extrusion mechanisms. We hypothesized that the chromatin is organized into spatially uncorrelated and segregated packing domains with various sizes and fractal dimensions. While some of the domains might be spatially isolated (domains 1,2,7,9), others may interpenetrate at the domain periphery as a result of decreasing chromatin density (domains 3 and 6, 5 and 8). For both configuration, the mass scaling outside of the domain is the same as the dimensionality of the space the chromatin is embedded in: 3.

We segmented the chromatin using automatic thresholding and performed the mass-scaling analysis (Fig. 3b). We calculated the chromatin mass enclosed by a ring with inner radius *r* and a single pixel width (discrete increment of the mass-scaling) as a function of *r* for different chromatin regions (Fig. 3c-d, red circles in (b)). From the average mass-scaling curve estimated from 4096 randomly centered chromatin regions (Fig. 3e), three regimes of the power-law scaling behavior can be identified: the chromatin chain region (yellow, r < 8.7 nm), the fractal region (blue, 8.7 nm < r < 50 nm, D = 2.6), and the space-filling region (region, r > 100 nm, D = 3). The transition between the fractal to the space filling region ranges from 50 to 100 nm (purple). This may indicate that at the supranucleosomal scale chromatin is organized into spatially uncorrelated (D > 3 at the supradomain scale) fractal packing domains with different sizes (100 - 200 nm in diameter) and fractal dimensions (D < 3 within domains) (Fig. 3f). While some domains are spatially isolated, others appear to be interpenetrating as a consequence of decreasing chromatin density at the domain periphery (Supplementary Fig.2).

### ChromSTEM analysis of nanoscale chromatin packing domains

We further investigated the local chromatin domain structure shown by the mass-scaling curve by calculating CVC, the average DNA concentration (Fig. 4a), and the variance of DNA concentration (Fig. 4b) in a 3D moving cube of 95.7 nm on each side and a stride of 2.9 nm. The CVC histogram shows (Fig. 4g) that the most probable CVC for A549 cells is 0.34, which is in good agreement with Ou’s measurements for an interphase human small airway cell (SEAC) ^24^. To estimate DNA concentration, we assumed that the highest voxel intensity in the tomograms corresponds to pure unhydrated DNA (~2 g/cm^3^) ^45^ and normalized all tomograms by the highest voxel intensity value. After normalization, DNA concentration 1 corresponds to DNA occupying 100% of the voxel volume or ~2 g/cm^3^. The average DNA concentration ranged from 0.006 to 0.38 with the most probable concentration at 0.14, and the variance of DNA concentration ranged from 0.0025 to 0.09 and peaked at 0.0342 (Fig. 4e, f).

**Figure 4.**
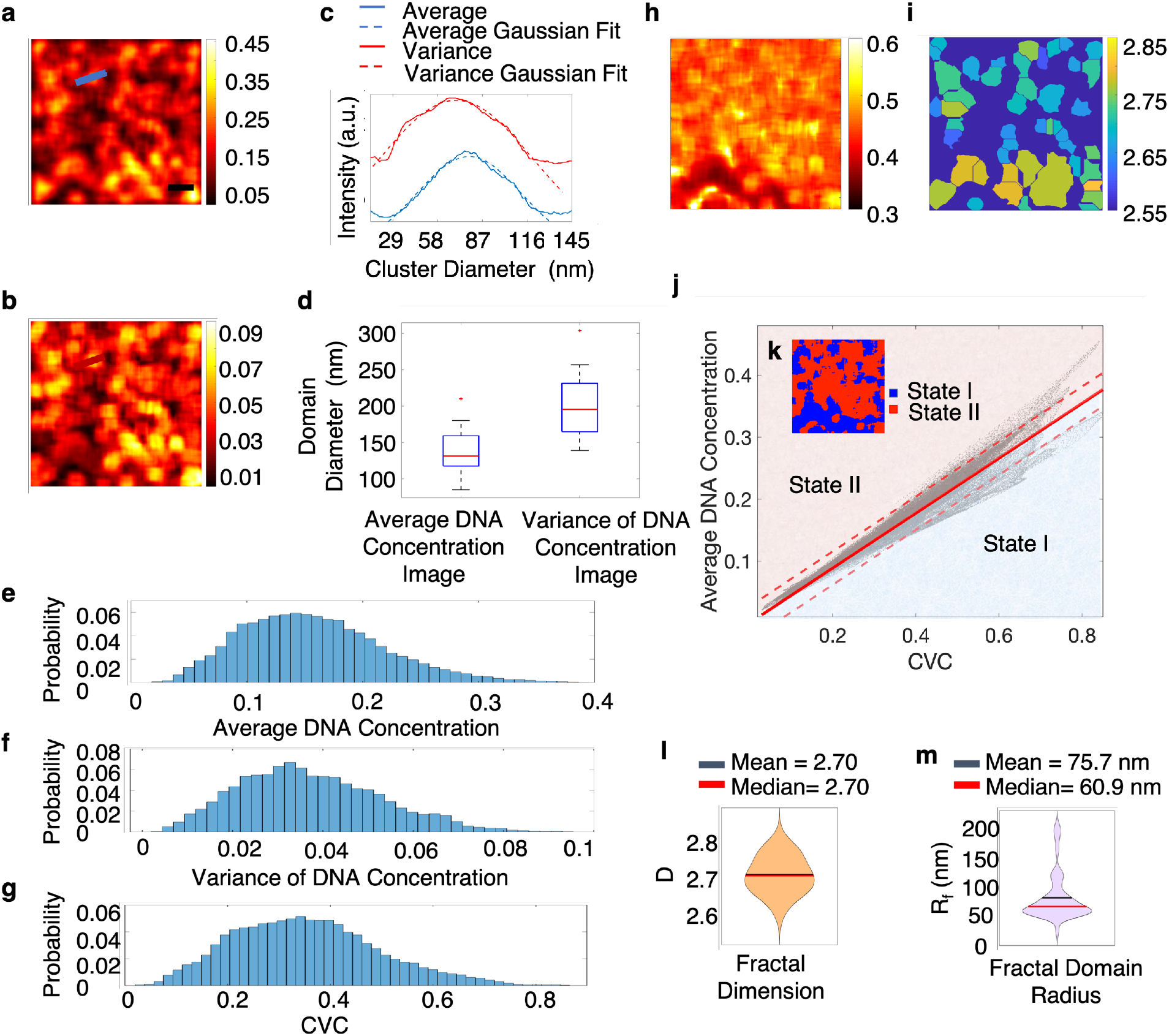
Chromatin is organized into spatially separated nanoscale packing domains. **(a)** The average DNA concentration and **(b)** the variance of DNA concentration both show chromatin packing domains. Scale bar: 100 nm for **(a,b,h,i)**. **(c)** The diameters of the domains were estimated as the full width at half maximum (HWHF) fitted from the line profile across the domain (blue line in **(a)** and red line in **(b)**). **(d)** Box plot of the domain diameter measured based on the maps of the average and the variance of DNA concentration. Histograms of **(e)** the average DNA concentration, **(f)** the variance of the DNA concentration, and **(g)** the CVC. **(h)** The map of the DNA content fraction defined as the ratio of the local average DNA concentration and the CVC. **(i)** The fractal dimension D for packing domains identified in **(a)**. **(j)** We identified two distinct states of chromatin packing with differential DNA content fraction (DNA rich vs. DNA poor domains). Red line: the linear regression of the entire dataset. DNA content fraction state I (blue) lies below the regression line, indicating low DNA fraction. State II (red) lies above the regression line, indicating high DNA fraction. The dashed line represents the 95% confidence interval of the linear regression. **(k)** Segmentation of chromatin based on DNA fraction. Chromatin regions with low DNA fraction (blue) and high DNA fraction (red) in **(k)** correspond to states I and II in **(j)**, respectively. From the mass-scaling curve for each of the 43 domains segmented by watershed algorithm from **(a)**, we quantified the fractal dimension **(l)** and fractal domain radius **(m)**.

The tomography data clearly shows a significant variability in local chromatin packing. In order to assess chromatin heterogeneity across the entire nucleus we analyzed ChromSTEM images of 30 nm cross sections of the whole nucleus of the same cell on which the tomography was performed. The average DNA concentration (Fig. 5b) ranged from 0.005 to 0.62, and the probability distribution showed a plateau of most probable DNA concentrations from 0.11 to 0.22. The variance of DNA concentration (Fig. 5d) ranged from 0.005 to 0.055 and peaked at 0.018.

**Figure 5.**
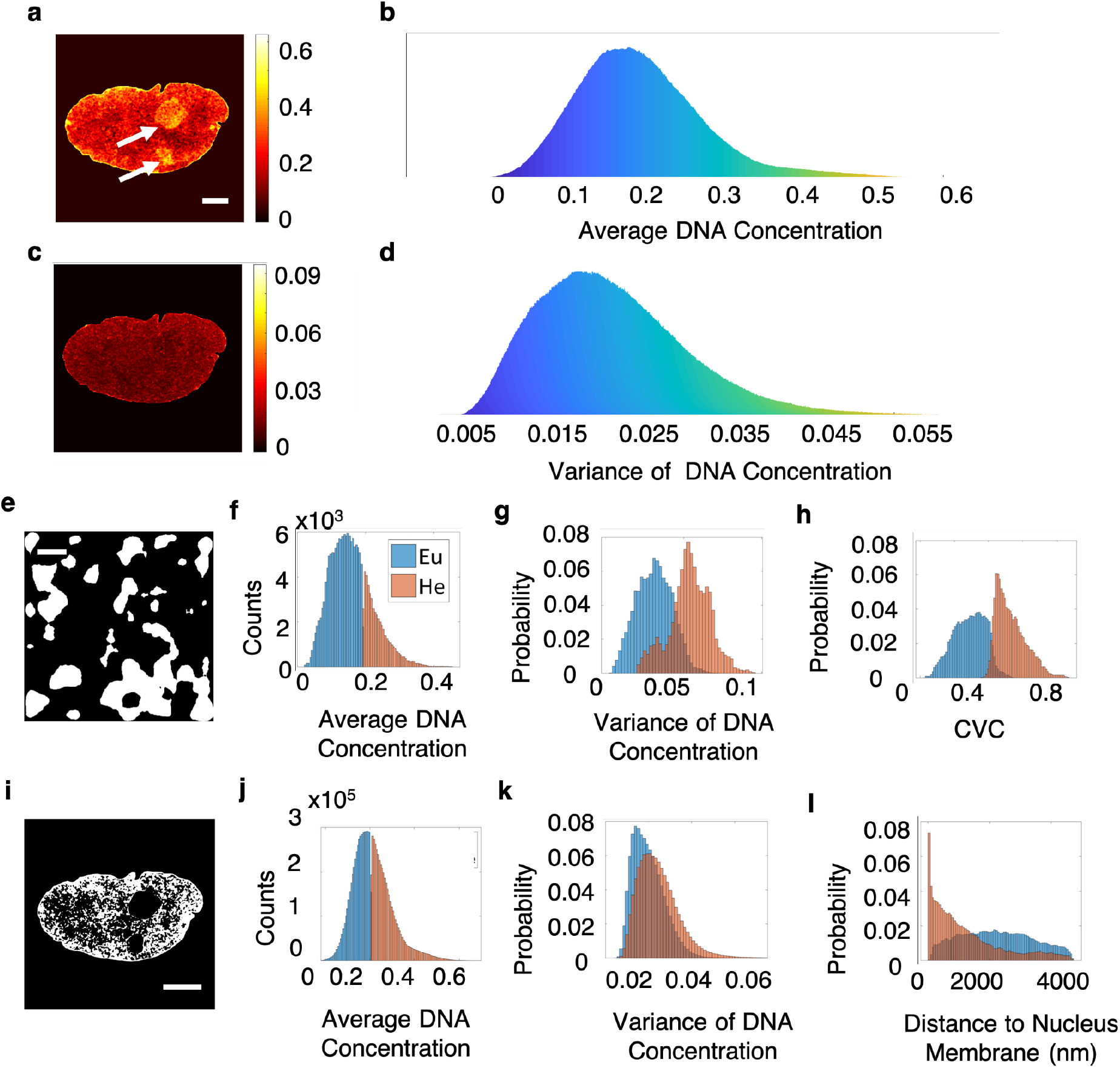
Packing properties of chromatin in different nuclear compartments. **(a)** The spatial and **(b)** the statistical distribution of the average DNA concentration calculated for a 30 nm section of A549 cells. Scale bar: 2 μm for **(a)** and **(c)**. The nucleoli are clearly identifiable (arrows in **(a)**). **(c)** The spatial and **(d)** the statistical distribution of the variance of the DNA concentration for the same sample. **(e)** Mask for heterochromatin segmentation based on the tomography data. Scale bar: 200 nm. **(f-h)** Differences in the average DNA concentration, the variance of the DNA concentration, and the CVC between euchromatin (blue) and heterochromatin (orange). **(i)** Mask for heterochromatin segmentation for the cell in **(a)**, the nucleoli were removed from the mask. Scale bar: 3 μm. **(j-l)** Differences in the average DNA concentration, the variance of the DNA concentration, and the distance to the nuclear envelope between euchromatin (blue) and heterochromatin (orange).

The spatial distribution of the domains with similar average DNA concentration (Fig. 4a) highly correlated with that of the domains with similar variance of the DNA concentration (Fig. 4b) (0.8 pixel-to-pixel Pearson’s correlation coefficient). The size of each domain was estimated as the full width at half maximum (FWHM) of the line profile across the domain (Fig. 4a blue line, 4b red line, 4c). Twenty-three domains were analyzed. The average diameter was 137.0 +/− 5.27 nm and 199.1 +/− 8.14 nm for domains defined based on their average DNA concentration and DNA concentration variance, respectively.

Notice that the highest CVC is 0.8 while the highest average DNA concentration is 0.4, indicating that the additional ~50% of the chromatin volume should be occupied by non-DNA molecules such as protein, which were not stained by chromEM methods. We then investigated the congruency of the CVC and the DNA concentration properties of chromatin in terms of DNA content fraction by dividing the average DNA concentration by CVC in the same moving cube (Fig. 4h). Linear regression was performed for data from all locations in the chromatin (Fig. 4j, regression line: solid red, 95% confidence interval: dashed red). As shown in Fig. 4K, the chromatin appears to fall into one of the two states: the DNA-poor (Region I) and the DNA-rich (Region II) based on whether the DNA content fraction is greater than the slope of the regression line. Nearly all packing domains could be classified into one of the two states (DNA-poor vs. DNA-rich), although the difference was more pronounced for domains with high average DNA concentration. Indeed, the boundaries of domains defined based on their average DNA concentration and the DNA content fraction were nearly identical (Supplementary Fig. 3). The mechanism behind such differential chromatin packing is unclear but might be related to phase separation ^46, 47^.

We further calculated the fractal dimension *D* (Fig. 4i, l) and the upper size of the domain fractality *R*_f_ (Fig. 4m) for packing domains segmented by watershed algorithm from the average DNA concentration map. *D* was obtained from the mass-scaling curve (*M*(*r*) ∝ *r*^*D*^), and *R*_f_ was estimated as the largest *r* for which the mass-scaling curve does not deviate from the power-law by more than 1%. Both mean and median *D* for all 43 domains was 2.7+/− 0.008. The mean and median *R*_f_ were 152.1 +/− 12.28 nm and 127.6 nm, respectively, in agreement with the domain size estimated from the average DNA concentration. In terms of the genomic size for the packing domains, given that the mean DNA concentration within a 95.7 nm^3^ cube is 0.157 (~ 0.314 g/cm^3^), the average molecular weight for single nucleotide is 325 Daltons, and the radius of the domain is 50 - 100 nm, we estimated that the packing domains contain 100 to 400 kb, comparable to the typical size of TADs ^48^.

### Comparison of chromatin packing for different nuclear compartments

Although it is frequently assumed that heterochromatin is denser compared to euchromatin, the difference in DNA density in these compartments is a subject of an ongoing debate. Some studies suggest that heterochromatin is significantly denser than euchromatin with implications that the transcriptional suppression in heterochromatin might in part be due to its higher density, which limits molecular diffusion processes. Other studies suggest that heterochromatin is only marginally denser than euchromatin ^49^. To provide quantitative data to address this question, we segmented heterochromatin by thresholding the average DNA concentration map from both the tomography data and the 30 nm cross sections of the nucleus (Fig. 4e, i). With this segmentation, the mean DNA concentration of heterochromatin (0.28) was approximately 2-fold higher than that of euchromatin (0.15) We also compared the variances of DNA concentration (Fig. 5g) and CVC (Fig. 5h) for the tomography data and the variance of DNA concentration (Fig. 5k) and the distance between euchromatin and the nuclear envelope (Fig. 5l) for the cross section data. The latter will provide a more complete view of the chromatin intra-cellular heterogeneity. In both cases, heterochromatin showed a larger mean variance than euchromatin. Regarding CVC, we observed a small overlap between heterochromatin and euchromatin regions. We also found that heterochromatin primarily resided along the inner nuclear membrane, with a small portion scattered across the whole nucleus: more than half of heterochromatin was adjacent to or within 500 nm from the nuclear envelope while only 7% of the euchromatin located within that range.

### Comparison of chromatin packing for different nuclear compartments in mouse ovary

It is becoming increasingly accepted that cell behavior *in situ* might in many instances be different from that in a cell culture, which may potentially apply to chromatin structure and function. We therefore demonstrated the ability of ChromSTEM to quantify chromatin structure in tissue samples. 40 μm mouse ovary tissue sections were labelled following the ChromEM protocol, and 120 nm thick resin sections were imaged with STEM HAADF contrast (Fig. 6a-e). Contrary to the findings in the A549 cells, we observed a clear differentiation in image contrast between euchromatin and heterochromatin, with the heterochromatin appearing significantly brighter than the euchromatin. The average DNA concentration was calculated for each cell and normalized to have the same range as the average DNA concentration of the A549 nucleus (Fig. 6f). The normalization did not influence the shape of the DNA concentration histograms. We calculated the probability distribution function of the average DNA concentration (Fig. 6f, red line), and the DNA concentration for euchromatin and heterochromatin (Fig. 6g). Of note, in all cells measured in the mouse ovary the probability density function of DNA concentration had a bimodal shape, as opposed to a single peak in case of the A549 cells. The most probable average DNA concentrations for euchromatin and heterochromatin were 0.195 and 0.468, respectively. Thus, DNA in heterochromatin can be almost 3-fold denser than that in euchromatin, and 2.4 times denser on average. This data indicates that the density of heterochromatin and euchromatin may have significant cell-to-cell variability even for the cells of the same type, suggesting the need to study chromatin organization across cell populations for accurate statistical conclusions.

**Figure 6.**
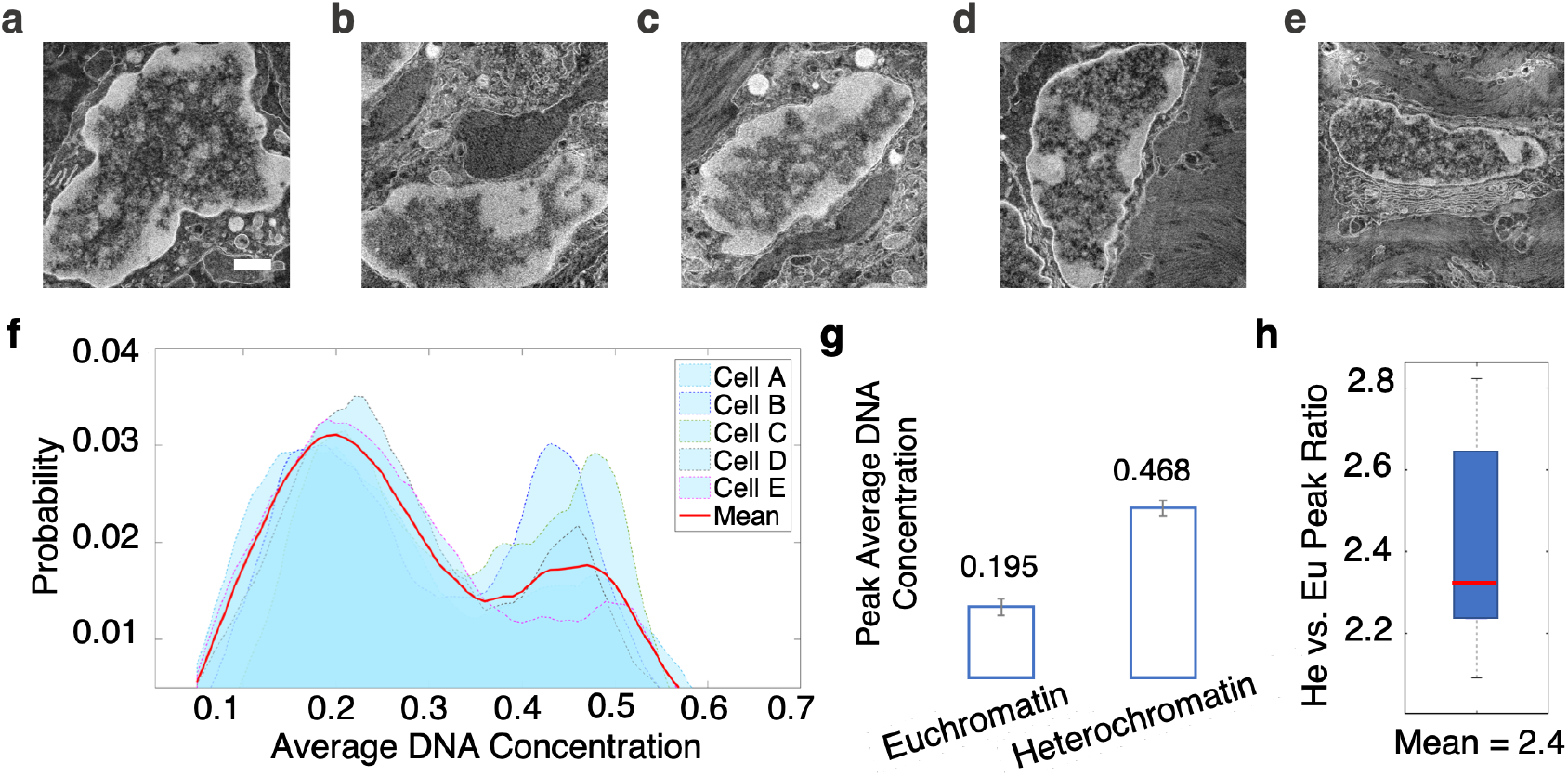
Differences in DNA concentration between euchromatin and heterochromatin of cell nuclei in the mouse ovary. **(a-e)** Neighboring cells in the mouse ovary tissue processed with the ChromEM staining. Scale bar: 1 μm. **(f)** The average DNA concentration for each cell (blue area) and the mean DNA concentration of the five cells (red line). **(g)** The most probable DNA concentration for euchromatin and heterochromatin. **(h)** The distribution of the ratio between the most probable DNA concentration within heterochromatin (He) and euchromatin (Eu) in each cell. On average, DNA concentration in heterochromatin is 2.4 times higher than that in euchromatin.

## Discussion

We developed a novel imaging platform to study 3D genome organization, ChromSTEM, by incorporating STEM tomography and imaging with Z-contrast into the original ChromEMT framework that provides contrast through DNA-specific staining. Importantly, obtaining the average and the variance of DNA concentration from the analysis of thin sections has miniscule error compared to the information gained from 3D tomography (Supplementary Fig. 4), which demonstrates the potential of the ChromSTEM platform for high-throughput chromatin analysis through thin section imaging. We demonstrated the versatility of ChromSTEM by quantifying the local chromatin packing structure and positioning through statistical metrics such as the scaling of the chromatin polymer, CVC, the average and the variance of DNA concentration, the DNA content fraction, and the localization of low- and high- density chromatin within the nucleus for cells *in vitro* as well as *ex vivo* tissue. Although the assumption that the ChromEM staining is stoichiometric does not hold at small length scales (single nucleotide) due to the diffusion inefficiencies of the dyes, in this work we utilized the binarized chromatin mask for the mass-scaling analysis. Moreover, the average and the variance of DNA concentration were estimated within a super-voxel. At the super-voxel level (100 nm on each side), we can assume the staining strength is uniform. Enhancing staining efficiency will improve the precision of ChromSTEM significantly at the small length scale and should be the focus of future research.

Utilizing ChromSTEM, we observed a “core-shell” structure of the chromatin chain (Fig. 2i) with the diameter comparable to that of the disordered polymer chain reported using ChromEMT by Ou et. al. ^24^ At the supranucleosomal level, we identified a high spatial heterogeneity of CVC that ranged from 6% to 74% and peaked at 34%, again, in agreement with the previous ChromEMT results. We also identified spatially distinct packing domains with 100 - 200 nm in diameter and quantified their internal CVC and the average and the variance of DNA concentration. Importantly, the domains could be further divided into two types based on their DNA content fraction: DNA rich vs. DNA poor. DNA poor domains are expected to have a substantially higher fraction of other molecular constituents such as proteins. Although these two alternate states of chromatin domains appear to exist in both euchromatin and heterochromatin, the difference between the two states were more pronounced in heterochromatin.

For most polymeric system, the 3D conformation can be described by several critical length scales: 1. The size of the basic chain size; 2. A 3D structure formed by the basic chain, which is typically characterized by a fractal (power-law) scaling; 3.The upper boundary of the fractal packing. While a polymer with uniform properties along its linear chain would typically form a single fractal conformation, polymers with linear properties that vary along their linear chain may form multiple fractal domains with distinct internal structure, in part driven by phase separation. As histone states, DNA methylation, loop formation, and DNA supercoiling may influence chromatin conformation, it is likely that chromatin is an example of the latter (Fig. 3f). We utilized ChromSTEM to elucidate the 3D conformation of chromatin across all these length scales and also to better understand the origin of the high heterogeneity of DNA density. We observed a three-regime power-law relationship between chromatin physical and genomic size (Fig. 3b): the chain region (r < 8.7 nm, D = 2.7), the supranucleosomal fractal domain (r above 8.7 nm and below 50 - 100 nm, D < 3, with D varying across domains with the average D = 2.6), and the space filling supradomain region (r > 100 nm, mass-scaling = 3). Both the length scale of transition from the fractal to the space filling region as well as D varied across domains, supporting the existence of multiple fractal chromatin structures. The average genomic size of the fractal domains was 100 to 400 kbp, comparable to the median size of a TAD ^48^. Similar values of D and domain sizes were reported by small angle neutron scattering and fluorescence correlation spectroscopy ^23, 50^. Models predict that such fractal structure imposes moderate diffusion hindrance by euchromatin, which is independent of the size of a diffusing molecule up to 100 nm diffusion range and should allow most biological macromolecules and even large macromolecular complexes access their targets within euchromatin ^50^. On the other hand, the high DNA concentration and CVC of heterochromatin is likely to significantly reduce chromatin accessibility, which may foster a transcriptionally silent state. This is also consistent with Molecular Brownian Dynamics simulations of transcription that have shown a dramatic suppression of diffusion of transcription factors in chromatin when its CVC exceeds 50% ^51^.

At the chromatin compartment level, our data indicates that heterochromatin has a substantially (more than 2-fold) higher DNA concentration compared to euchromatin, which potentially addresses a perplexing question of the differences in density between euchromatin and heterochromatin with some of the prior studies suggesting that euchromatin is considerably denser than euchromatin while others arguing that the two compartments may have comparable density ^19, 50, 52^. In future studies, precise segmentation of chromatin compartments based on labeling histone markers using correlative 3D optical super resolution and electron microscopy will need to be performed to further improve the measurement of chromatin compartments. Future studies may also elucidate whether the two types of DNA rich vs. DNA poor domains correspond to functionally distinct states of chromatin such as the gene rich or gene poor subtypes.

## Methods

### Cell culture and tissue biopsy collection

The A549 adenocarcinoma human lung epithelial cells were grown to reach confluency of 60% in DMEM with 10% FBS and 1x penicillin/streptomycin in 35 mm MatTek dishes (MatTek Corp) at 37°C at 5% CO_2_. Animal procedures were performed at NorthShore University Health System, with the approval of the Institutional Animal Care and Use Committee (IACUC). A healthy Fisher 344 rat (150–200 g; Harlan) was euthanized and the biopsy was harvested from the ovary and immersed in EM fixative immediately at room temperature for 20 min then transferred to fresh fixative and stored at 4°C overnight.

### ChromEM sample preparation

The A549 cells were prepared using the ChromEM method published previously, and lists of reagents and step-by-step protocols can be found in the supplementary information. The biopsy was embedded in low melting point agarose (Thermo Fisher) and 40 μm thick sections were prepared using a vibratome (VT1200 S Leica) on ice. The sections were deposited onto a glass-bottom petri-dish (MatTek Corp) and treated as cultured cells for ChromEM preparation.

For photo-oxidation, an inverted microscope (Eclipse, Nikon Inc.) with a 100x objective with LED lamp was employed. A cold stage was developed in-house from a wet chamber equipped with humidity and temperature control. For all analysis in the paper, we performed photo-oxidation for 7 min for each region, and fresh DAB solution was used for every time. We investigated the influence of photo-oxidation time by illumination two regions in the same dish for 7 min and 25 min respectively (Supplementary Fig. 5).

100 nm thick sections of a A549 cell were made by ultramicrotomy (UC7, Leica) and deposited onto a plasma-treated slot grid with carbon/Formvar film (EMS). Then 10 nm colloidal gold nanoparticles were deposited on both sides as fiducial markers for ChromSTEM tomography. Ultrathin sections with a nominal thickness of 30 nm for A549 cells and 40 nm for tissue were made and deposited onto a plasma-treated mesh grid with Formvar/carbon film (EMS) for ChromSTEM imaging.

### EM data collection and tomography reconstruction

A 200kV STEM (HD2300, HITACHI) with HAADF mode was employed for all image collection. For tomography, the sample was tilted from −60° to 60° with 2° increments on two roughly perpendicular axes. The pixel size was chosen to be 2.9 nm to resolve the fiducial markers. Each tilt series was aligned with fiducial markers in IMOD and reconstructed using Tomopy with penalized maximum likelihood for 40 iterations independently. IMOD was used to combine the tomograms to suppress artifacts (Supplementary Fig. 1), the nominal voxel size is 2.9 nm. The ultrathin sections were imaged at 0 tilt angle with a pixel size of 5.4 nm.

### Mass-scaling analysis

In polymer physics, the mass-scaling is the relationship between the material *M* within concentric circles of radius *r* and the radius *r*. For a fractal structure, the mass scales as *M*(*r*) ∝ *r*^*D*^, where *D* is the power-scaling exponent, or fractal dimension. In reality, the fractal structure can be altered by the physical chemical environment, and the critical length scale where the mass-scaling deviates from power law is defined as the fractal domain radius. Importantly, the mass-scaling is translational invariant, regardless of the choice of the center of the concentric circles. Prior to mass-scaling analysis, the chromatin position was segmented by automatic thresholding with Li’s method in FIJI as previous reported ^24^. For each scaling analysis, concentric circles with radius from 2.9 nm (1 pixel) to 290 nm (100 pixels) were employed, and non-zero pixels were chosen as the origin of the mass-scaling on the binarized chromatin mask. As the stack is only 100 nm thick, we employed mass-scaling on each virtual 2D slice then averaged through z direction to get average mass-scaling for local region (Fig. 3c-d). For the entire tomogram, 4096 centers were randomly selected on each of the 33 slices, the average mass-scaling curve was used in quantifying fractal dimension (D) and the transition length scale (Fig. 3e).

### Calculating average, variance of DNA concentration and CVC

As STEM HAADF contrast originates from Rutherford scattering, the tomography intensity is proportional to and can be converted to the physical mass of DNA. We normalized the tomogram to its highest intensity to extract the DNA concentration in each voxel. We assumed the highest voxel intensity corresponded to unhydrated DNA (2g/cm^3^), or the voxel was permeated only with DNA. Then the new voxel intensity corresponds to the fraction of the voxel is DNA (0.5 mean 50% space is occupied by DNA with a density of 1g/cm^3^). Then we employed a moving 3D cube (95.7 nm in each dimension and stride of 2.9 nm) to analyze the local chromatin packing. We mapped the mean and the variance of DNA concentration within the cube on the normalized tomogram (Fig. 4a-b, e-f). We adopted the definition of CVC from previous work ^24^, and obtained the CVC using the same cube dimension on the binarized chromatin mask by calculating the fraction of non-zero voxels in each cube (Fig. 4g).

For ultrathin sections, the moving cube (square with thickness) was anisotropic, the height of the cube was limited by the section thickness. Nonetheless, the lateral dimension of the square was chosen such that the volume stayed the same as the one used in tomography data: we used 170 nm wide square for 30 nm sections and 147 nm for 40 nm sections. We calculate the average and the variance of DNA concentration from normalized STEM projection images and proofed that no or minimal error were introduced compared to true 3D analysis (Supplementary Fig. 4). To render more physical value, we first estimated the range of average DNA concentration for the whole nucleus on the thick section with tomography data (Supplementary Fig. 6) and scaled the average DNA concentration from ultrathin sections to have the same range (Fig. 5a-b, 6f), we employed the same pre-factor in rescaling the variance of DNA concentration (Fig.5 c-d).

### Chromatin Packing Domain Analysis

#### Domain Size

From tomography data, we identified nanoscale packing domains with similar average or variance of DNA concentration. To estimate the size of the domains, we manually selected 23 domains and calculated the full-width-half-maximum of the line profile across the domain (Fig 4c). To obtain the fractal dimension of each domain, we first segment the domains from the DNA concentration map using intensity thresholding and watershed algorithm.

#### Domain fractal dimension and radius from mass-scaling analysis

Similar to mass-scaling analysis for the entire field of view, for each domain, we calculated the average mass-scaling curve centered on every non-zero pixel within the domain for each slice in the binarized 2D chromatin mask. We quantified the average fractal dimension by calculating the power law exponent on the average mass-scaling curve, and coded every pixel using the same fractal dimension within that domain (Fig. 4i). We estimated the fractal domain radius by finding the spatial separation with 1% deviation from the power-law scaling on the average mass-scaling curve. Violin plot was employed to show the distribution of fractal dimension and fractal domain radius for 44 domains (Fig 4l-m).

#### DNA fraction analysis

The CVC is the fraction of space chromatin occupied, and the average DNA concentration is the fraction of space DNA occupied. We divided the average DNA concentration by the CVC for the same moving cub to calculate the fraction of DNA on chromatin (Fig. 4h). We plotted the average DNA concentration vs. CVC for each pixel and performed linear regression considering all data points to separate two packing states (Fig. 4j): DNA-poor (above the regression line) and DNA-poor (below the regression line).

#### Segmenting different nuclear compartments

We quantified the heterochromatin volume percentage within the A549 cell nucleus using STORM (Supplementary Fig.7). On average, the heterochromatin accounts for 47% of the total chromatin. Assuming the heterochromatin is denser than the euchromatin, we calculated that the threshold of average DNA concentration that provided a 47%-53% split for A549 cells was 0.2. Regions with average DNA concentration above 0.2 were considered heterochromatin and the rest euchromatin. For tissue samples, the histogram of the average DNA concentration showed two peaks, the peak position was employed to represent the most probably average DNA concentration for different chromatin compartments.

## Supporting information

Supplementary Mov. 6

Supplementary Mov. 5

Supplementary Mov. 4

Supplementary Mov. 3

Supplementary Mov. 2

Supplementary Mov. 1

## Acknowledgement

This work was supported by National Institutes of Health grants U54CA193419, R01 CA228272, R01CA225002 and National Science Foundation grants EFMA-1830961 and EFMA-1830969. The authors also thank the BioCryo and EPIC facilities of Northwestern University’s NUANCE Center, which has received support from the Soft and Hybrid Nanotechnology Experimental (SHyNE) Resource (NSF ECCS-1542205); the MRSEC program (NSF DMR-1720139) at the Materials Research Center; the International Institute for Nanotechnology (IIN); the Keck Foundation; and the State of Illinois, through the IIN. The imaging methodology is partially based on research sponsored by the Air Force Research laboratory under agreement number is FA8650-15-2-5518. The U.S. Government is authorized to reproduce and distribute reprints for Governmental purposes notwithstanding any copyright notation thereon. The views and conclusions contained herein are those of the authors and should not be interpreted as necessarily representing the official policies or endorsements, either expressed or implied, of Air Force Research Laboratory or the U.S. Government.

## Author Contributions

Y.L. designed experiments, conducted the ChromEM sample preparation, the STEM tomography and imaging, the data analysis, and prepared the manuscript. E.R., V.A. assisted the ChromEM sample preparation. A.E. and J.F. performed STORM sample preparation and imaging. L.A., R.B. contributed ideas for experiments and manuscript. A.S. contributed data analysis. V.P.D., V.B. directed and supervised the project. All authors discussed the results and contributed to the manuscript.

## Reference

1. Yu M, Ren B. The three-dimensional organization of mammalian genomes. Annual review of cell and developmental biology 33, 265–289 (2017).

2. Clowney EJ, et al. Nuclear aggregation of olfactory receptor genes governs their monogenic expression. Cell 151, 724–737 (2012).

3. Bickmore WA. The spatial organization of the human genome. Annual review of genomics and human genetics 14, 67–84 (2013).

4. Gorkin DU, Leung D, Ren B. The 3D genome in transcriptional regulation and pluripotency. Cell stem cell 14, 762–775 (2014).

5. van Steensel B, Belmont AS. Lamina-associated domains: links with chromosome architecture, heterochromatin, and gene repression. Cell 169, 780–791 (2017).

6. Dekker J, Mirny L. The 3D genome as moderator of chromosomal communication. Cell 164, 1110–1121 (2016).

7. Tang Z, et al. CTCF-mediated human 3D genome architecture reveals chromatin topology for transcription. Cell 163, 1611–1627 (2015).

8. Almassalha LM, et al. Macrogenomic engineering via modulation of the scaling of chromatin packing density. Nature biomedical engineering 1, 902 (2017).

9. Li G, et al. Extensive promoter-centered chromatin interactions provide a topological basis for transcription regulation. Cell 148, 84–98 (2012).

10. Wang H, et al. CRISPR-Mediated Programmable 3D Genome Positioning and Nuclear Organization. Cell 175, 1405–1417. e1414 (2018).

11. Montavon T, et al. A regulatory archipelago controls Hox genes transcription in digits. Cell 147, 1132–1145 (2011).

12. Furlan-Magaril M, Várnai C, Nagano T, Fraser P. 3D genome architecture from populations to single cells. Current opinion in genetics & development 31, 36–41 (2015).

13. de Wit E, et al. The pluripotent genome in three dimensions is shaped around pluripotency factors. Nature 501, 227 (2013).

14. Iwafuchi-Doi M, Zaret KS. Pioneer transcription factors in cell reprogramming. Genes & development 28, 2679–2692 (2014).

15. Zhang Z, Pugh BF. High-resolution genome-wide mapping of the primary structure of chromatin. Cell 144, 175–186 (2011).

16. Gibcus JH, Dekker J. The hierarchy of the 3D genome. Molecular cell 49, 773–782 (2013).

17. Rao SS, et al. A 3D map of the human genome at kilobase resolution reveals principles of chromatin looping. Cell 159, 1665–1680 (2014).

18. Nagano T, et al. Single-cell Hi-C reveals cell-to-cell variability in chromosome structure. Nature 502, 59 (2013).

19. Le Gros MA, et al. Soft X-ray tomography reveals gradual chromatin compaction and reorganization during neurogenesis in vivo. Cell reports 17, 2125–2136 (2016).

20. Shi G, Liu L, Hyeon C, Thirumalai D. Interphase human chromosome exhibits out of equilibrium glassy dynamics. Nature communications 9, 3161 (2018).

21. Szabo Q, et al. TADs are 3D structural units of higher-order chromosome organization in Drosophila. Science Advances 4, eaar8082 (2018).

22. Boettiger AN, et al. Super-resolution imaging reveals distinct chromatin folding for different epigenetic states. Nature 529, 418 (2016).

23. Lebedev D, et al. Fractal nature of chromatin organization in interphase chicken erythrocyte nuclei: DNA structure exhibits biphasic fractal properties. FEBS letters 579, 1465–1468 (2005).

24. Ou HD, Phan S, Deerinck TJ, Thor A, Ellisman MH, O’shea CC. ChromEMT: Visualizing 3D chromatin structure and compaction in interphase and mitotic cells. Science 357, eaag0025 (2017).

25. Li Y, et al. Measuring the Autocorrelation Function of Nanoscale Three-Dimensional Density Distribution in Individual Cells Using Scanning Transmission Electron Microscopy, Atomic Force Microscopy, and a New Deconvolution Algorithm. Microscopy and Microanalysis 23, 661–667 (2017).

26. Schwarzer W, et al. Two independent modes of chromatin organization revealed by cohesin removal. Nature 551, 51 (2017).

27. Almassalha LM, et al. The global relationship between chromatin physical topology, fractal structure, and gene expression. Scientific reports 7, 41061 (2017).

28. Almassalha LM, et al. The greater genomic landscape: the heterogeneous evolution of cancer. Cancer research, (2016).

29. Solovei I, et al. Nuclear architecture of rod photoreceptor cells adapts to vision in mammalian evolution. Cell 137, 356–368 (2009).

30. Gürsoy D, De Carlo F, Xiao X, Jacobsen C. TomoPy: a framework for the analysis of synchrotron tomographic data. Journal of synchrotron radiation 21, 1188–1193 (2014).

31. Kremer JR, Mastronarde DN, McIntosh JR. Computer visualization of three-dimensional image data using IMOD. Journal of structural biology 116, 71–76 (1996).

32. Mastronarde DN. Dual-axis tomography: an approach with alignment methods that preserve resolution. Journal of structural biology 120, 343–352 (1997).

33. Schindelin J, et al. Fiji: an open-source platform for biological-image analysis. Nature methods 9, 676 (2012).

34. Acemel RD, Maeso I, Gómez-Skarmeta JL. Topologically associated domains: a successful scaffold for the evolution of gene regulation in animals. Wiley Interdisciplinary Reviews: Developmental Biology 6, e265 (2017).

35. Narendra V, Bulajić M, Dekker J, Mazzoni EO, Reinberg D. CTCF-mediated topological boundaries during development foster appropriate gene regulation. Genes & development 30, 2657–2662 (2016).

36. Cremer T, Cremer C. Chromosome territories, nuclear architecture and gene regulation in mammalian cells. Nature reviews genetics 2, 292 (2001).

37. Dekker J, Heard E. Structural and functional diversity of topologically associating domains. FEBS letters 589, 2877–2884 (2015).

38. Sexton T, Cavalli G. The role of chromosome domains in shaping the functional genome. Cell 160, 1049–1059 (2015).

39. Dixon JR, Gorkin DU, Ren B. Chromatin domains: the unit of chromosome organization. Molecular cell 62, 668–680 (2016).

40. Sexton T, et al. Three-dimensional folding and functional organization principles of the Drosophila genome. Cell 148, 458–472 (2012).

41. Dixon JR, et al. Topological domains in mammalian genomes identified by analysis of chromatin interactions. Nature 485, 376 (2012).

42. Bintu B, et al. Super-resolution chromatin tracing reveals domains and cooperative interactions in single cells. Science 362, eaau1783 (2018).

43. Huang K, Backman V, Szleifer I. Interphase chromatin as a self-returning random walk: Can DNA fold into liquid trees? bioRxiv, 413872 (2018).

44. Stevens TJ, et al. 3D structures of individual mammalian genomes studied by single-cell Hi-C. Nature 544, 59 (2017).

45. Panijpan B. The buoyant density of DNA and the G+ C content. Journal of chemical education 54, 172 (1977).

46. Larson AG, Narlikar GJ. The role of phase separation in heterochromatin formation, function, and regulation. Biochemistry 57, 2540–2548 (2018).

47. Erdel F, Rippe K. Formation of chromatin subcompartments by phase separation. Biophysical journal, (2018).

48. Hansen AS, Cattoglio C, Darzacq X, Tjian R. Recent evidence that TADs and chromatin loops are dynamic structures. Nucleus 9, 20–32 (2018).

49. Huisinga KL, Brower-Toland B, Elgin SC. The contradictory definitions of heterochromatin: transcription and silencing. Chromosoma 115, 110–122 (2006).

50. Bancaud A, Huet S, Daigle N, Mozziconacci J, Beaudouin J, Ellenberg J. Molecular crowding affects diffusion and binding of nuclear proteins in heterochromatin and reveals the fractal organization of chromatin. The EMBO journal 28, 3785–3798 (2009).

51. Matsuda H, Putzel GG, Backman V, Szleifer I. Macromolecular crowding as a regulator of gene transcription. Biophysical journal 106, 1801–1810 (2014).

52. Imai R, et al. Density imaging of heterochromatin in live cells using orientation-independent-DIC microscopy. Molecular biology of the cell 28, 3349–3359 (2017).

