## Supplementary Information for "Quantifying Three-dimensional Chromatin Organization Utilizing Scanning Transmission Electron Microscopy: ChromSTEM"

**Supporting Information**

### Methods

#### ChromEM sample preparation

The ChromSTEM sample staining and resin-embedding followed the published protocol, and detailed reagents and steps can be found in S.I. All cells were thoroughly rinsed in Hank's balanced salt solution without calcium and magnesium (EMS) before fixation with EM fixative. Two stages of fixation were performed: room temperature fixation for 5 min and on-ice fixation for an hour with fresh fixative. The cells were kept cold for all following steps before resin embedding either on ice or on a cold stage with the temperature monitored to vary from 4 $^{o}$C to 10 $^{o}$C. The biopsy of the mouse ovary was embedded in low melting point agarose (Thermo Fisher) and 40 µm thick sections were prepared using a vibratome (VT1200 S Leica) on ice. The sections were deposited onto a glass-bottom petri-dish (MatTek) and treated as described for cells in the following steps.

After fixation, the samples were bathed in blocking buffer for 15 min before being stained by DRAQ5$^{TM}$ (Thermo Fisher) for 10 min. The cells were rinsed and kept in blocking buffer before photo-bleaching and submerged in 3-5'-diaminobenzydine (DAB) solution (Sigma Aldrich) during photo-bleaching on the cold stage.

A Nikon microscope (Nikon Inc.) was used for photo-bleaching. A cold stage was developed in-house from a wet chamber equipped with humidity and temperature control. The influence of illumination intensity was studied by applying different photo-bleaching time for two different spots in the same dish: 7 min and 25 min, respectively (S1 Fig). For the study of chromatin packing, the photo-bleaching time for each spot was 7 min, and two spots were illuminated in each dish.

After photo-bleaching, the cells were rinsed in 0.1M sodium cacodylate buffer thoroughly. Reduced osmium solution (EMS) was used to enhance the contrast in STEM HAADF mode, and the heavy metal staining lasted 30 min on ice. Serial ethanol dehydration was performed, and during the last 100% ethanol wash, the cells were brought back to room temperature. Durcupan resin (EMS) was used for embedding after infiltration, and the blocks were cured at 60$^{o}$C for 48 hrs.

An ultramicrotome (UC7, Leica) was employed to prepare sections of different thicknesses. For STEM HAADF tomography, 100 nm thick sections of a A549 cell were made and deposited onto a copper slot grid with carbon/Formvar film. 10 nm colloidal gold fiducial markers were deposited on both sides of the sample. For STEM HAADF projection imaging, ultrathin sections with a nominal thickness of 30 nm of A549 cells were made and deposited onto a copper mesh grid with Formvar/carbon coating (EMS). All TEM grids were plasma cleaned prior to sectioning and no post-staining was performed to the sections.

**Supplementary Movie 1.** First tilt series for the STEM-HAADF tomography.

**Supplementary Movie 2.** Second tilt series for the STEM-HAADF tomography.

**Supplementary Movie 3.** Tomogram of 3D chromatin structure moving through z direction.

**Supplementary Movie 4.** 3D rendering of the entire tomography stack.

**Supplementary Movie 5.** 3D rendering of nucleosomal superstructure and linker DNAs.

**S1 Table. Reagents used in ChromSTEM staining**

| Reagent | Formula |
|---|---|
| Washing solution | Hank's balanced salt solution without calcium and magnesium |
| Fixation solution | 2.5% EM grade glutaraldehyde<br>2% paraformaldehyde<br>2 mM $CaCl_2$<br>0.1 M sodium cacodylate buffer, pH = 7.4 |
| Blocking solution | 10 mM glycine<br>10 mM potassium cyanide<br>0.1 M sodium cacodylate buffer, pH = 7.4 |
| DNA staining solution | 10 µM DRAQ5<br>0.1% SAPONIN<br>0.1 M sodium cacodylate buffer, pH = 7.4 |
| Bathing solution | 2.5 mM 3,3'- diaminobenzidine tetrahydrochloride (DAB)<br>0.1 M sodium cacodylate buffer, pH = 7.4 |
| Reduced osmium staining solution | 2% osmium tetroxide<br>1.5% potassium ferrocyanide<br>2 mM $CaCl_2$<br>0.15 M sodium cacodylate buffer, pH = 7.4 |
| Durcupan™ resin mixture 1 | 10 mL Durcupan ™ ACM single component A, M, epoxy resin<br>10 mL Durucupan ™ ACM single component B, hardener 964<br>0.15 mL Durcupan ™ ACM single component D |
| Ducrupan ™ resin mixture 2 | 10 mL Durcupan ™ ACM single component A, M, epoxy resin<br>10 mL Durucupan ™ ACM single component B, hardener 964<br>0.2 mL Durcupan ™ ACM, single component C, accelerator 960<br>0.15 mL Durcupan ™ ACM single component D |
| 1:1 infiltration mixture | 10 mL 100% ethanol<br>10 mL Durcupan ™ resin mixture 1 |
| 2:1 infiltration mixture | 5 mL 100% ethanol<br>10 mL Durcupan ™ resin mixture 1 |

**Supplementary Protocol 1. Sample preparation for ChromSTEM for cell cultures**

**Fixation:**

1. Wash the cells in the petri-dish in the washing solution for 3 times, 2 minutes each.
2. Fix the cells with the fixation solution for 5 minutes at room temperature.
3. Continue to fix the cells with fresh fixation solution for an additional 1 hour on ice.

The following steps before the last ethanol dehydration are either on ice or on a cold stage, all reagents must be chilled to 4$^{o}$C prior to use.

**DNA Staining:**

4. Wash the cells with 0.1M sodium cacodylate buffer for 5 times on ice, 2 minutes each.
5. Block the cells with blocking solution for 15 minutes.
6. Stain the cells with DNA staining solution for 10 minutes.
7. Wash the cells with the blocking solution for 3 times, 5 minutes each.

**Photo-bleaching:**

8. Bath the cells in the bathing solution prior to photo-bleaching
9. Photo-bleach the cells using continuous epi-fluorescence illumination (150 W Xenon Lamp) with Cy5 red tilter and a 100x objective for 7 minutes for each spot on the cold stage.
10. Replace the bathing solution in the petri-dish with fresh bathing solution every 15 minutes (roughly two spots).

**Heavy metal staining:**

11. Rinse the cells with 0.1 M sodium cacodylate buffer for 5 times, 2 minutes each.
12. Stain the cells with reduced osmium staining solution for 30 minutes.
13. Wash the cells with double distilled water for 5 times, 2 minutes each.

Dehydration and Resin embedding

14. Dehydrate the cells with serial ethanol (30%, 50%, 70%, 85%, 95%, 100% twice) on ice, 2 minutes each.
15. Wash the cells with 100% ethanol at room temperature for 2 minutes.
16. Infiltrate the cells with 1:1 infiltration mixture at room temperature for 30 minutes.
17. Infiltrate the cells with 2:1 infiltration mixture at room temperature for 2 hours.
18. Infiltrate the cells with Durcupan™ resin mixture 1 at room temperature for 1 hour.
19. Infiltrate the cells with Durcupan™ resin mixture 2 at 50 $^{o}$C in the dry oven for 1 hour.
20. Flat embed the cells with fresh Durcupan™ resin mixture 2 in Beem capsule and cure at 60 $^{o}$C in the dry oven for 48 hours.

**Supplementary Protocol 2. Sample preparation for ChromSTEM for tissue biopsies**

**Harvest mouse ovary (Performed by PDX tumor core in Northwestern University)**

1. Euthanize the mice and harvest the ovary
2. Dissect the ovary with ideal orientation using a double-edged platinum knife to smaller pieces (~$4^3$ $mm^3$ cubes).
3. Store the tissue biopsies on ice-cold 0.1 M PBS for no more than 30 minutes before the sectioning.

**Section the tissue with vibratome**

4. Embed the tissue cubes in 5% low temperature agarose and chill at $4^oC$ for 5 minutes
5. Section the agarose-tissue sample in ice-cold 0.1 M PBS using a vibratome (Leica) to 40 µm (roughly 3 layers of cells) slices.
6. Deposit the thin slices onto a plasma treated glass-bottom petri-dish (MatTek) coated with poly-L-lysine, and cover with ice-cold 0.1 M PBS. The tissue slice will adhere to the bottom of the dish due to gravity and poly-L-lysine. DO NO leave the tissue in PBS for more than 5 minutes before chemical fixation.

**ChromSTEM tissue sample preparation**

7. Treat the tissue slice in the petri-dish as the cell culture and repeat S1 Protocol for the rest of ChromSTEM sample preparation.

#### Supplementary Protocol 3. Sample preparation and Imaging for Photon localization Microscopy of A549

##### Photon Localization Microscopy sample preparation

A549 lung adenocarcinoma cells were cultured on 35 mm glass bottom dishes until approximately 70% confluent. Cells were washed with phosphate-buffered saline (PBS) for 2 minutes then fixed with a solution of 3% Paraformaldehyde and 0.1% Glutaraldehyde in PBS for 10 minutes. Cells were washed for 5 minutes in PBS, then quenched in 0.1% sodium borohydride in PBS for 7 minutes. Cells were washed 3 times in PBS for 5 minutes each, then permeabilized in blocking buffer (0.2% Triton X-100 and 3% Bovine serum albumin (BSA) in PBS) for 20 minutes. The primary antibodies to target heterochromatin (anti-H3K9me3, Abcam and anti-H3K27me3, Abcam) was added to the blocking buffer to a concentration 2.5 µg/mL and incubated for 2 hours. Cells were then washed in washing buffer (0.1% Triton X-100 and 0.2% BSA in PBS) 3 times for 5 minutes. Cells were then incubated with the secondary antibody (Alexa Fluor 647, Thermo Fisher Scientific) at a concentration of 2.5 µg/mL in blocking buffer for 40 minutes. Cells were then washed two time in PBS for 5 minutes each. Cells were imaged in standard imaging buffer with an oxygen scavenging system containing 0.5 mg/mL glucose oxidase (Sigma-Aldrich), 40 µg/mL catalase (Roche or Sigma-Aldrich), 143 mM 2-hydroxy-1-ethanethiol, and 100 mg/mL glucose in TN buffer (50 mM Tris (pH 8.0) and 10 mM NaCl)

##### Photon Localization Microscopy Imaging

Photon Localization Microscopy (PLM) was used to quantify the fraction of chromatin that is heterochromatin. Samples were prepared according to protocols in Sec. XX. Samples were imaged on an inverted microscope base (Eclipse Ti-U with perfect-focus system, Nikon). Samples were illuminated with a 637 nm laser (Obis, Coherent), which was passed through a bandpass filter (FF01-637/7-25, Semrock), reflected off a dichroic beam splitter (Di01-R405/488/532/635-25x36, Semrock) and sent through a 100X 1.49 NA objective (SR APO TIRF, Nikon) with an average power at the sample of 3 to 15kW per $cm^3$. Fluorescence emission images were collected via the 100X objectives and projected through a notch filter (#67-120, Edmund Optics) to reject any reflected laser light and onto an EMCCD (iXon Ultra 888, Andor). Multiple frames of stochastic “blinking” events were captures and super resolution PLM images were reconstructed using the ThunderSTORM plugin.

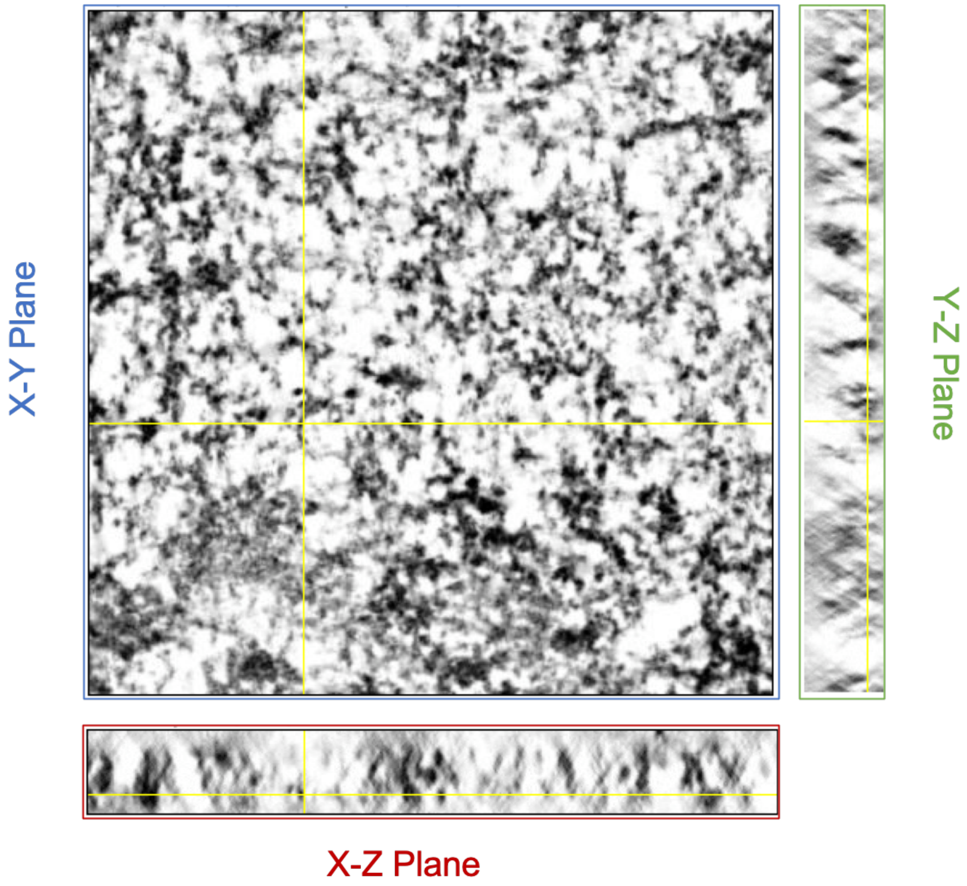


**Supplementary Figure 1.** Orthogonal views of chromatin after tomography reconstruction of a A549 cell. In the tomography experiment and reconstruction, dual-tilt and penalized maximum likelihood algorithm were employed to suppress the artifacts introduced by the missing cone. In the X-Z and Y-Z plane, the “X” shaped artifacts can still be seen in the tomograms, but the individual nucleosomes can be easily identified, and the stretch in the Z direction is not severe. The quality of this tomography is not as high as the ones used in Ou’s ChromEMT work, as only two axes were used in our work but eight in his work. However, based on the CVC analysis, our tomography exhibits an almost identical histogram as the one shown in Ou’s paper, indicating our tomography has sufficient quality for studying the local chromatin packing for 100 nm super-voxels.

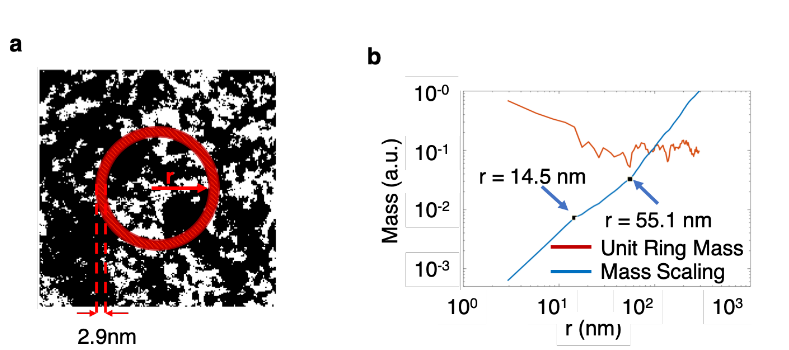


**Supplementary Figure 2.** Mass-scaling and density scaling. **(a)** The mass scaling of the fractal region was calculated as described. The density distribution was the unit mass on the ring with inner radius r and bandwidth of 2.9 nm (1 pixel). **(b)** The mass scaling and unit ring mass (density) for the mask in (a). Two transitions are present. In r<14.5 region, the density decreases following a power law. In 14.5 nm < r < 55.1 nm, the density keeps dropping but with oscillations but overall following the same scaling as expected from a mass fractal. At r = 55.1 nm, the density increases sharply. In r>55.1 nm, the density oscillates but maintains the baseline. This behavior indicates interpenetrating of the domains.

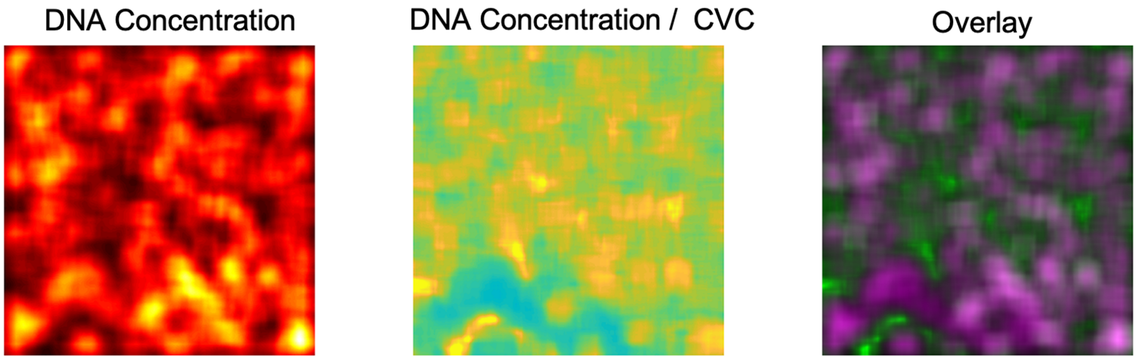


**Supplementary Figure 3.** Comparing cluster boundaries in the DNA concentration and the DNA concentration / CVC. To investigate the spatial distribution of cluster identified independently in the DNA concentration map (left) and the DNA concentration/CVC ratio map (middle), we overlaid the two maps with false coloring (right). In the overlay image, the magenta denotes the DNA concentration, and the green represents the DNA concentration/CVC ratio. Qualitatively, the clusters have similar boundaries and spatial distribution. However, the cluster with high DNA concentration can have arbitrary value of DNA concentration/CVC ratio, which leads to low pixel-to-pixel cross correlation coefficient (-0.13).

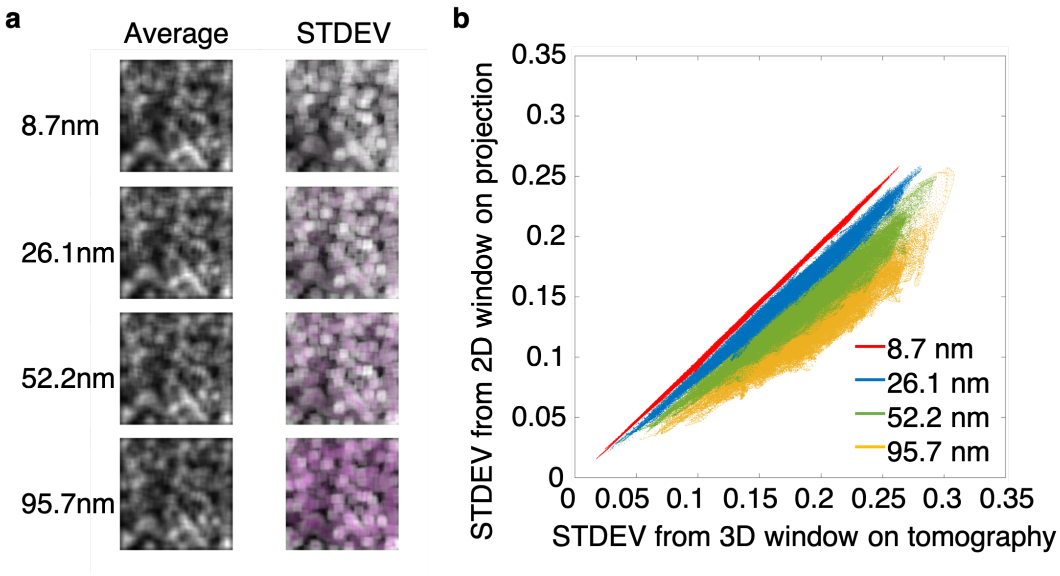


**Supplementary Figure 4.** Influence of projection at different thickness to the average and the standard deviation of the DNA concentration. We projected various numbers of the virtual 2D slices after tomography reconstruction to form "projections" and calculated the 2D average and the 2D standard deviation (STDEV) of DNA concentration using a 2D window with 95.7 nm on each side. For each projection (thickness), the 3D average and the 3D STDEV were calculated using a 3D window with 95.7 nm in x and y and the thickness of the projection in Z. In theory, the 2D and the 3D average of DNA concentration should remain the same, while the 2D STDEV will be a non-monotonic underestimation of the 3D STDEV. **(a)** We coded the 2D metrics magenta and the 3D metric green and overplayed them for different thickness. For each pixel, a perfect match of the two will result in black and white contrast, a mismatch will result in colored contrast. As expected, for all thickness, the average from 2D and 3D window matched perfectly. While for STDEV, the mismatch increases rapidly as the thickness of projection increases. **(b)** The STDEV calculated from 2D window on the projection was plotted against the STDEV calculated from 3D window on the tomography to quantify the extent of underestimation given this chromatin structure at different thickness. From 8.7 nm to 95.7 nm, the ratio of 2D STDEV to 3D STDEV kept decreasing, indicating at larger thickness, the STDEV is more severely underestimated. Meanwhile, the spread of the curve at larger thickness is also significantly wider, indicating a non-monotonic, irreversible smearing of the STDEV. However, at small thickness (ultra-thin section) such as 8.7 nm, the slope is 1 and the spread is minimal, indicating almost no smearing of the STDEV. For 26.1 nm projection, the slope is 0.9 with a $r^2$ equals to 0.99 in the linear regression, suggesting that the STDEV from the projection is sufficiently accurate to serve as the proxy of the real STDEV from tomography with a pre-factor difference.

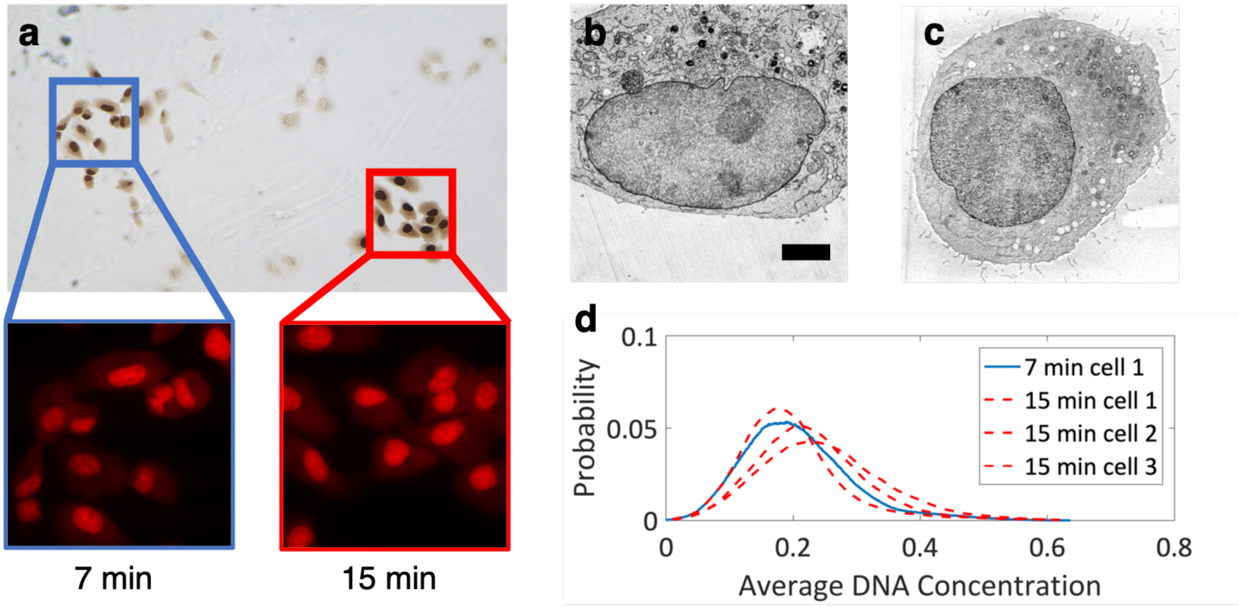


**Supplementary Figure 5.** Difference in staining A549 cell by varying photo-bleaching time. To test the consistency of photo-bleaching, we experimented the process with different illumination time. **(a)** Photobleaching for 7 min (blue square) and for 15 min (red square). The resulting staining is significantly heavier in the long photo-bleaching spot. **(b)** STEM HAADF image (contrast inverted) of the 30 nm section for one cell in the 7 min spot. Scale bar: 2 µm. **(c)** TEM image of the 30 nm section for one cell in the 15 min spot. For the TEM images, we first converted the image contrast to mass-thickness using the Beer's lambert law, then calculated the average DNA concentration, and normalized the histogram to the same range as the STEM image. **(d)** Comparison of the histogram of the average DNA concentration for one cell in the 7 min spot (blue solid line) and three cells in the 15 min spot (red dash lines). We observed that the average DNA concentration of the 7 min cell lied in the range of the 15 min cells. Considering cell to cell variations, we concluded that within the time frame, there is no significant influence of the length of photo-bleaching in the analysis of the chromatin packing.

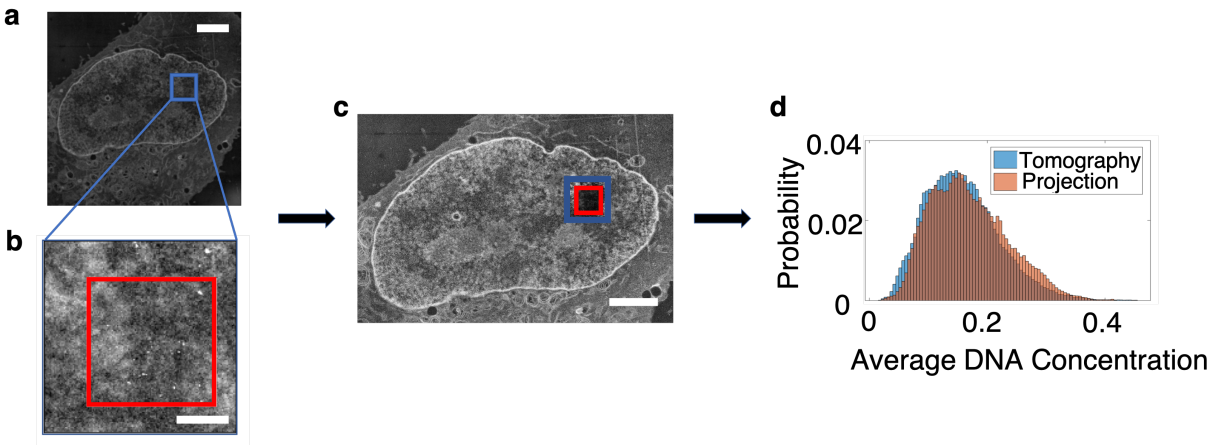


**Supplementary Figure 6.** Alignment of the tomography region to the whole nucleus and the normalization of average DNA concentration of A549 cells. To quantify the DNA concentration for the entire nucleus, the tomography region was registered to the whole nucleus using cross correlation with openCV2 packing in Python, and the average DNA concentration calculated from the whole nucleus image was normalized to the average DNA concentration calculated directly from the tomography. **(a)** A larger ROI (blue square) including the tomography region was selected for the automatic registration. Scale bar: 2 µm. **(b)** The tomography region (red square) was registered to the ROI. Scale bar: 500 nm. **(c)** Overlay the tomography region (red square) and the ROI (blue square) onto the whole nucleus image. Scale bar: 2 µm. The histogram of the average DNA concentration of the tomography region but calculated from the whole cell projection (orange) was normalized to the average DNA concentration calculated from the tomography (blue). The coefficients were used to normalize the DNA concentration for the whole nucleus. We observed small discrepancies for the average DNA concentration calculated from the tomography and projection even after normalization, we believe the difference in noise level at two image condition might be the reason. Importantly, the majority of the histograms match perfectly.

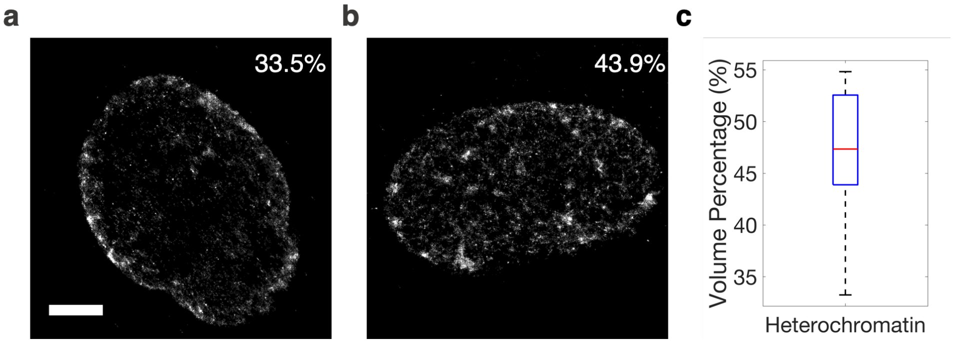


**Supplementary Figure 7.** STORM of A549 to quantify heterochromatin volume concentration. To quantify the average volume percentage of heterochromatin for A549 cell, the H3K9me3 and anti-H3K27me3 were labeled and STORM images were taken for multiple cells. The ratio of pixels with signal and total pixels of the nucleus was used to represent the average heterochromatin volume percentage. **(a)** and **(b)** Examples of STORM images with 33.5% and 43.9% heterochromatin, scale bar: 3μm. **(c)** Distribution of heterochromatin volume percentage for 4 cells at in total 10 different focal planes, the average heterochromatin volume concentration is calculated to be 47% and used in heterochromatin segmentation from average DNA concentration map.
